# TMEM106B in humans and Vac7 and Tag1 in yeast are predicted to be lipid transfer proteins

**DOI:** 10.1101/2021.03.12.435176

**Authors:** Tim P. Levine

## Abstract

TMEM106B is an integral membrane protein of late endosomes and lysosomes involved in neuronal function, its over-expression being associated with familial frontotemporal lobar degeneration, and under-expression linked to hypomyelination. It has also been identified in multiple screens for host proteins required for productive SARS-CoV2 infection. Because standard approaches to understand TMEM106B at the sequence level find no homology to other proteins, it has remained a protein of unknown function. Here, the standard tool PSI-BLAST was used in a non-standard way to show that the lumenal portion of TMEM106B is a member of the LEA-2 domain superfamily. The non-standard tools (HMMER, HHpred and trRosetta) extended this to predict two yeast LEA-2 proteins in the lumenal domains of the degradative vacuole, equivalent to the lysosome: one in Vac7, a regulator of PI(3,5)P_2_ production, and three in Tag1 which signals to terminate autophagy. Further analysis of previously unreported LEA-2 structures indicated that LEA-2 domains have a long, conserved lipid binding groove. This implies that TMEM106B, Vac7 and Tag1 may all be lipid transfer proteins in the lumen of late endocytic organelles.

## Introduction

Proteins of unknown function persist as a sizable minority in all organisms, with 15% of yeast and human proteins still having no informative description of their function at the molecular level.^1^ Even if mutation or deletion of a protein links its function to a specific cellular pathway, the direct action of the protein might be at some distance from the observed pathway.^2^ TMEM106B (previously called FLJ44732) is a type II transmembrane protein named in a generic way because its function was not obvious from its sequence.^3^ Interest in TMEM106B first arose when the gene was linked with familial frontotemporal lobar degeneration with TDP-43 inclusions.^4,5^ A parallel genetic link was found in Alzheimer’s Disease with the same inclusions.^6^ While these phenotypes result from overexpression of TMEM106B, a different neuronal phenotype, demyelination, is found both with D252N mutation,^7,8^ and with deletion in an animal model.^9^ Outside the brain, raised TMEM106B drives metastasis of K-Ras-positive lung cancer.^10,11^ With attention turning to coronavirus biology since the SARS-CoV-2 pandemic, TMEM106B has been repeatedly identified as a protein required to support productive SARS-CoV-2 infection.^12–14^

In cell biological studies, the TMEM106B protein has been localised to late endosomes and lysosomes,^3,11,15,16^ and it has been shown to be important for many lysosomal functions, including: maintaining normal lyosomal size,^15–18^ net anterograde transport of lysosomes along axons,^17,18^ and transcriptional programmes that upregulate lysosomal components,^11^ including those required for acidification.^15,19^ Thus, over-production of lysosomal proteases might explain its role in cancer metastasis.^10,11^ Homologues of TMEM106B have only previously been described in animals. Humans are typical of chordates in expressing three homologues, with TME016B accompanied by two unstudied but closely related paralogues (TMEM106A/C), all between 250-275 residues.^3^ In comparison, invertebrates tend to either have one homologue, or none – for example TMEM106 is missing from all insects. The cytoplasmic N-terminus of TMEM106B (residues 1-96) is unstructured.^20^ Following a single transmembrane helix (TMH), there is a lumenal C-terminal domain of 157 residues, which has five glycosylation sites.^3^

Beyond localisation and topology, studies of TMEM106B have made limited progress. Other than that the D252N mutation causes hypomyelination,^7^ which mimics loss of function,^9^ no structural information is available, either experimental or predicted. A major factor that might have contributed to TMEM106B remaining among the proteins of unknown function at the molecular level is that no homologues are available for comparative cell biological study in genetically tractable model organisms, including *Drosophila, C. elegans*, and both budding and fission yeast.^21^ To address this, we examined the sequence of TMEM106B using bioinformatics tools. The standard tool PSI-BLAST was used in a non-standard way to show that the C-terminal intra-lysosomal domain of TMEM106B, its most conserved portion, belongs to the little studied but widely spread LEA-2 domain superfamily. Next, two yeast LEA-2 homologues were identified: Vac7, a regulator of PI(3,5)P_2_ generation, and Tag1 a regulator of autophagy. TMEM106B, Vac7 and Tag1 localisations are all lysosomal (equivalent to degradative vacuole in yeast). The homology is greatest between TMEM106B and Vac7, where the TMHs show sequence similarity. Examination of the previously unreported solved structure of an archaeal LEA-2 domain showed that it is a lipid transfer protein, which suggests specific modes of action for TMEM106B, Vac7 and Tag1, along with all LEA-2 proteins, related to sensing and/or transferring lipids.

## Methods

### Structural Classification of Proteins at SUPERFAMILIES

Standard searches were carried out with all proteins of interest at the SUPERFAMILY database.^22^

### Conservation Analysis

Protein conservation for TMEM106B was assessed by creating a representative multiple sequence alignment (MSA) in four steps: (i) gathering DUF1356 sequences (PFAM07092) from the full set of representative proteomes (n=838);^23^ (ii) clustering using MMSeq2 with default settings and reducing each cluster with to a single member (n=238);^24^ (iii) aligning these with MUSCLE^25^, which outperforms other MSA tools;^26^ (iv) removing short sequences (here <150 aa) or those with deletions in key conserved regions, suggesting splicing errors. This left 208 sequences. JALVIEW was used to extract conservation scores from the alignment.^27^ The same pathway was followed for 3BUT.

### Domain Composition

Domain composition in proteins returned by PSI-BLAST (Supplementary Tables 1 and 5) was determined by searching annotations both in name and domain fields. Accepted alternative terms for LEA-2 domains were: nonrace-specific disease resistance-1 (NDR1), Harpin-induced (HIN), and yellow-leaf-specific gene-9 (YLS9). Remaining unassigned sequences were submitted to the National Library of Medicine’s Conserved Domains Database search tool.^28^ Non-significant hits in PSI-BLAST (Supplementary Table 5), refer to matches with E-values between 0.001 and 1.

The distribution of proteins across different fungal clades were determined from databases as followed: from PFAM – using Tree visualisations on Species Distribution tabs; from Uniprot and NCBI – combining domain search terms with fungal clade terms (Ascomycota, Basidiomycota, Mucoromycota, Zoopagomycota, Chytridiomycota and Blastocladiomycota). Membrane topologies were assessed with TMHMM 2.0 and Signal 5.0.^29,30^

### PSI-BLAST strategies

Initial, standard PSI-BLAST with TMEM106B (human) used the non-redundant database at NCBI (threshold E-value 0.001).^31^

PSI-BLAST to find more diverse hits for TMEM106B, LEA-2 protein, C-terminus of Vac7, Tag1 was performed at the Tuebingen Toolkit using a “nr50” version of NCBI database, which has been filtered so that the maximum pairwise sequence identity is 50%.^32^ The LEA-2 protein chosen as seed was an archaeal tandem LEA-2 protein (*Thermococcus litoralis*, WP_148290494.1, 311 aa). This is the typical size and form of archaeal LEA-2 proteins. Residues 185-309 are the closest known homologue to the sequence crystalised as 3BUT (125 residues align with E-value 5×10^-35^). The *T. litoralis* sequence was used to seed searches rather than 3BUT because the latter is fragment from the C-terminus of an *Archaeoglobus fulgidus* protein for which no complete sequence is in the database, only the incomplete sequence KUJ92443.1 (271 aa) being available.

### Iterative searching with Jackhmmer

Iterative searches building profiles with hidden Markov models were carried out in Jackhmmer, part of the HMMER suite using standard settings, *i.e*. cut-off E-values of 0.01 for the whole sequence and 0.03 for each hit.^33^

### Remote homology search with HHpred

HHpred was carried out using standard settings, (3 iterations of HHblits, Alignment Mode: no re-align) except the cut-off for multiple sequence alignment (MSA) generation was set Evalue ≤0.01. MSAs were forwarded back to HHpred to indicate areas of high homology by switching Alignment Mode to Realign with MAC, with Re-alignment Threshold set to 0.3 (default). In some instances, Re-alignment Threshold was set to 0.01 to extend alignment towards the ends of the query and target, even though the additional aligned areas did not add any statistical significance. Alignment to LEA-2 in HHpred was assessed from hits to the solved structure 3BUT in its database of solved structures. The Vac7 sequence submitted was the 287 residues between 879-1165 and variants missing either one or both regions 995-1036 and 1079-1118.

### Cluster Map

Sequences in six protein families were accumulated from HHblits searches (8 rounds, searching into UniRef30 pre-clustered database. These seeds, with resulting numbers of hits in brackets, were: Vac7 (454), TMEM106B (1489), DUF3712 protein (W9WCQ9 in *Cladophialophora*) (776), Tag1 (531), and two negative controls that showed some but not all characteristics of LEA-2 domains: DUF2393 (O25031 in *Helicobacter*) (645) and DUF3426 (Q9HUW2 in *Pseudomonas)* (753). These 4648 sequences were reconciled for repeats, by filtering to reduce similarity using MMseq2 with default settings,^24^ and the LEA-2 like domain was extracted as the 50 residues before the C-terminus of the TMH and at minimum 40 residues after, retaining a maximum of 240 residues after the TMH (or the first 290 residues if no TMH was identified). This produced 2810 sequences, the origins of which were: 794 TMEM106B only, 115 Vac7 only, 242 DUF3712 only, 192 Tag1, 189 DUF2393 only, 171 DUF3426 only, 245 in TMEM106B+Vac7, 346 TMEM106B+DUF3712, 2 in Vac7+DUF3712, 82 in TMEM106B+DUF3712+Vac7 and 432 in DUF2393+DUF3426. Sequences were blasted all vs all in CLANS.^34,35^ Clustering in 2D was carried out with default settings, with p-value threshold for inclusion set to 2×10^-4^, which excluded all DUF2393 or DUF3426 proteins (n=792) and 67 of the other 2018 proteins. Relationships between groups were repeatable. Links, in particular involving TMEM106B, Vac7 and DUF3712 were checked by hand for relevance.

### Structure Prediction

3D models of Vac7 and Tag1 were made by Phyre2 (intensive mode) and SWISS-MODEL(standard settings).^36,37^ Models of both TMEM106B and Vac7 made by analysis of contact co-evolution were made in trRosetta, switched either to ignore known structures or to use them as templates.^38^ 3D alignment of models with those already solved was carried out by the DALI server,^39^ performing either pairwise comparisons structure against the subset of structures in the Protein Data Bank where sequences are non-redundant at the 25% level (PDB25), or comparisons across a bespoke grid.

### Structure Visualisation

Structures were visualised using CCP4MG software. For NMR structures of LEA-2 domains 1YYC and 1xO8, a single structure was constructed with every atom in the average position of the 20 models provided. Surface colouring was either by electrostatic potential using the yellow red blue (YRB) scheme ^40^, or by conservation (scale blue➔red➔white, see key in Figure 4B).

## Results

### TMEM106B is the animal representative of the late embryogenesis abundant-2 (LEA-2) superfamily

To predict the protein fold of TMEM106B, we started with the Structural Classification of Proteins (SCOP) tool hosted at the Superfamily server.^22^ This predicted that the lumenal region of TMEM106B, a region that contains multiple conserved blocks, is in the superfamily of late embryogenesis abundant (LEA)-2 domains (E-value = 1×10^-5^) (Figure 1). This finding parallels an automated low confidence prediction in MODBASE (made in 2008, retrieved 2021).^41^ The LEA-2 domain superfamily (96 residues), alternatively named LEA14 or WHy (for upregulation in <u>W</u>ater stress and <u>Hy</u>persensitive response),^42^ has previously been reported to have members widely spread across bacteria, archaea and plants, but not in animals or fungi.^43^ Genes in the family share an overall phenotype of supporting cellular responses to stresses such as desiccation,^44^ but no molecular function has been described.^45,46^

**Figure 1:**
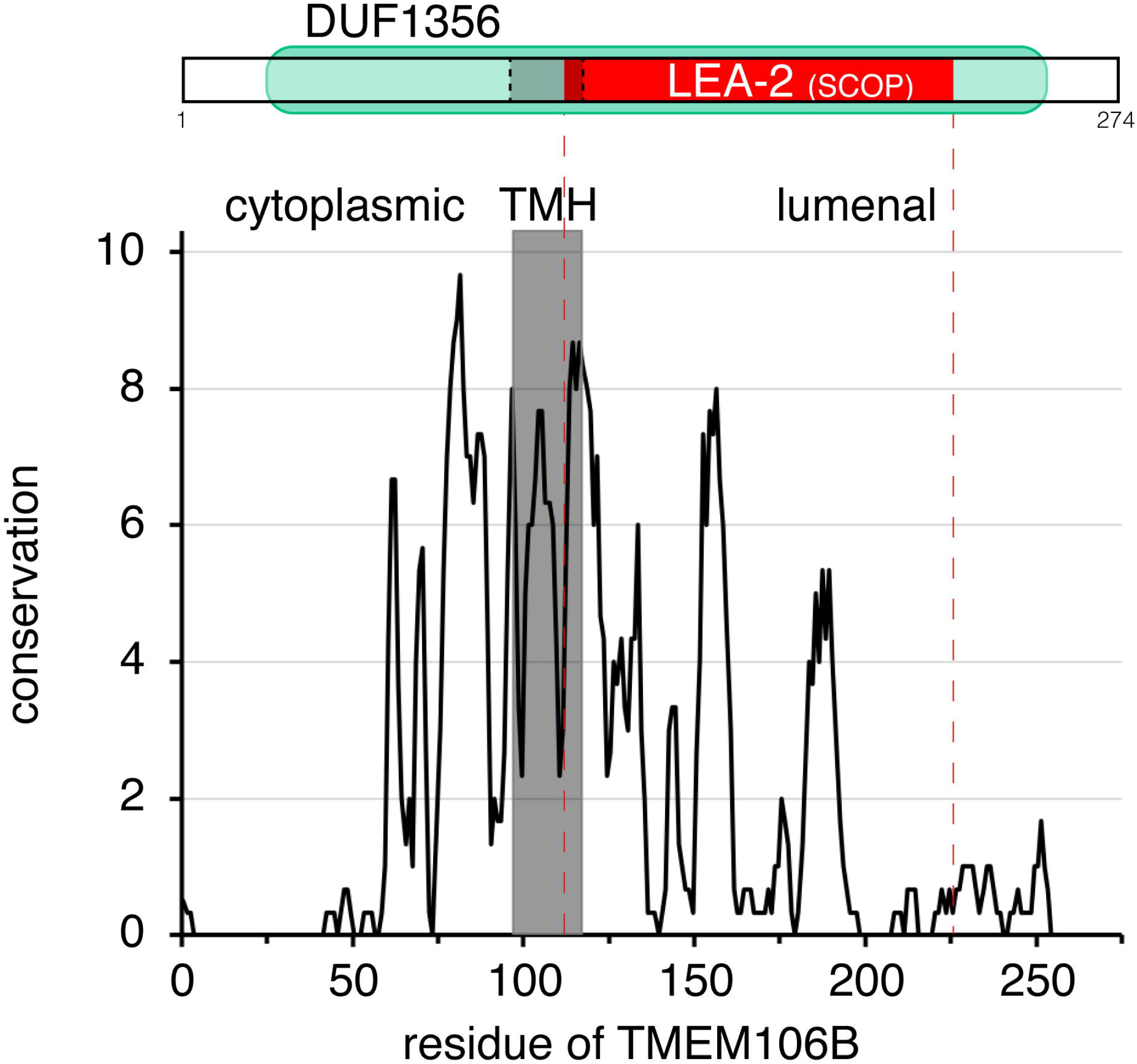
Conserved portions of the lumenal domain of TMEM106B are identified as homologous to LEA-2. Top: The Structural Classification of Proteins (SCOP) tool identified a region of homology in TMEM106B to LEA-2 (red) that includes the end of the transmembrane helix (TMH, gray with dashed borders) and much of the lumenal portion of DUF1356 (pale green). Bottom: Conservation across TMEM106B. Scores of 10 indicate all physico-chemical properties are conserved, and 11 indicates identity. Dashed red lines indicate limit of homology identified by SCOP.

To confirm the link between TMEM106B and LEA-2, we carried out detailed PSI-BLAST searches. Searching in the non-redundant NCBI database containing all sequences (nr100), the first iteration identified >2000 TMEM106B homologues, almost all in animals, and the iterative search converged rapidly thereafter (Supplementary Table 1). This result matches the distribution of TMEM106B both in the literature,^47^ and in the Protein Families (PFAM) database, which defines the central 80% of TMEM106B as the domain of unknown function-1356 (DUF1356, 228 residues, Figure 1), of which 99% are in animals and 1% in algae.^23^ An important feature of the nr100 database is that it is dominated by vertebrate sequences that are very close to the human seed,^47^ so these dominate the profile generated, leading the multiple sequence alignment (MSA) of hits to overly focus on the seed. Here, searching in nr100 likely prevented non-vertebrate sequences from diversifying the MSA. Therefore, we repeated the PSI-BLAST using a database pre-filtered so that the maximum pairwise sequence identity is 50% (nr50).^32^ The first iteration identified almost only known TMEM106B homologues, as with nr100 searches. However nr50 search differed from nr100 from the second iteration onwards by including LEA-2 hits, which increased in number and eventually dominated (Supplementary Table 1). Thus, a PSI-BLAST strategy that focusses on sequence diversity rather than allowing dominance by vertebrate sequences shows that TMEM106B is a sequence homologue of LEA-2, indicating that TMEM106B is in the LEA-2 superfamily.

### Vac7 is a predicted fungal member of the LEA-2 superfamily

While TMEM106B represent LEA-2 superfamily members in animals, this still leaves LEA-2 domains undocumented in fungi.^43^ To investigate this, the initial step was to examine databases of fungal proteins for automatically generated annotations as TMEM106B or LEA-2. The NCBI database, the largest numerically, has 436 fungal proteins annotated as LEA-2 homologues and 18 as TMEM106B (*i.e*. DUF1356, Supplementary Table 2A). Many fungal phyla are represented, except Ascomycota, the largest fungal phylum that includes the model organisms *S. cerevisiae* and *S. pombe* (Supplementary Table 2B).

We hypothesised that homologues within the LEA-2 superfamily may exist in Ascomycota, but that they have diverged below the level of detectability by PSI-BLAST, which has a limit of approximately 20-35% sequence identity.^48^ To find such remote homologues for both TMEM106B and LEA-2 we used two tools that are more sensitive than PSI-BLAST. The first approach was the HMMER Suite, which gains sensitivity over PSI-BLAST by using hidden Markov models to flexibly interpret profiles using bespoke rules, for example for gap penalties.^49^ It also limits searches to representative proteomes, avoiding domination by highly sequence clades of organisms. A profile built from TMEM106B using Jackhmmer had hits annotated as LEA-2 from the first iteration, and they dominated from iteration 3 (Supplementary Table 2A). From iteration 4 onwards an increasing number of hits were annotated as being homologues of the *S. cerevisiae* type II vacuolar membrane protein Vac7, named for its role in <u>VAC</u>uolar morphology, the fungal equivalent of the lysosome.^50^ The search aligned the hydrophobic region and the N-terminal 50 residues of the LEA-2 domain of TMEM106B with the same region in Vac7. In the reverse search, Vac7 linked to LEA-2 from iteration 2 onwards (Supplementary Table 2B). This shows that the vac7 protein family is also within the LEA-2 superfamily.

Similar results were obtained with HHsearch, a remote homology profile-profile tool that explicitly aligns predicted secondary structure,^51^ which is enacted on the HHpred online server.^32^ Searches seeded either with the C-terminus of TMEM106B or with a LEA-2 domain produced strong hits to each other, with probabilities that they are homologous of 98/99% and E-values for the alignment based on sequence alone of 2×10^-4^/3×10^-7^ (Figure 2A/B). With TMEM106B as seed, the top hit was the solved archaeal LEA-2 structure 3BUT (113 residues/127), which is the C-terminus of a type II membrane protein with tandemly repeated LEA-2 domains.^52^ The hit covered six β strands and one α helix and lacked strand 1 at the N-terminus of the domain (Supplementary File 1). Both LEA-2 domains of the archaeal protein produced reverse hits for TMEM106B, the N-terminal domain including the hydrophobic region and a domain of seven β strands plus one α helix (Figure 2B, Supplementary File 1).

**Figure 2:**
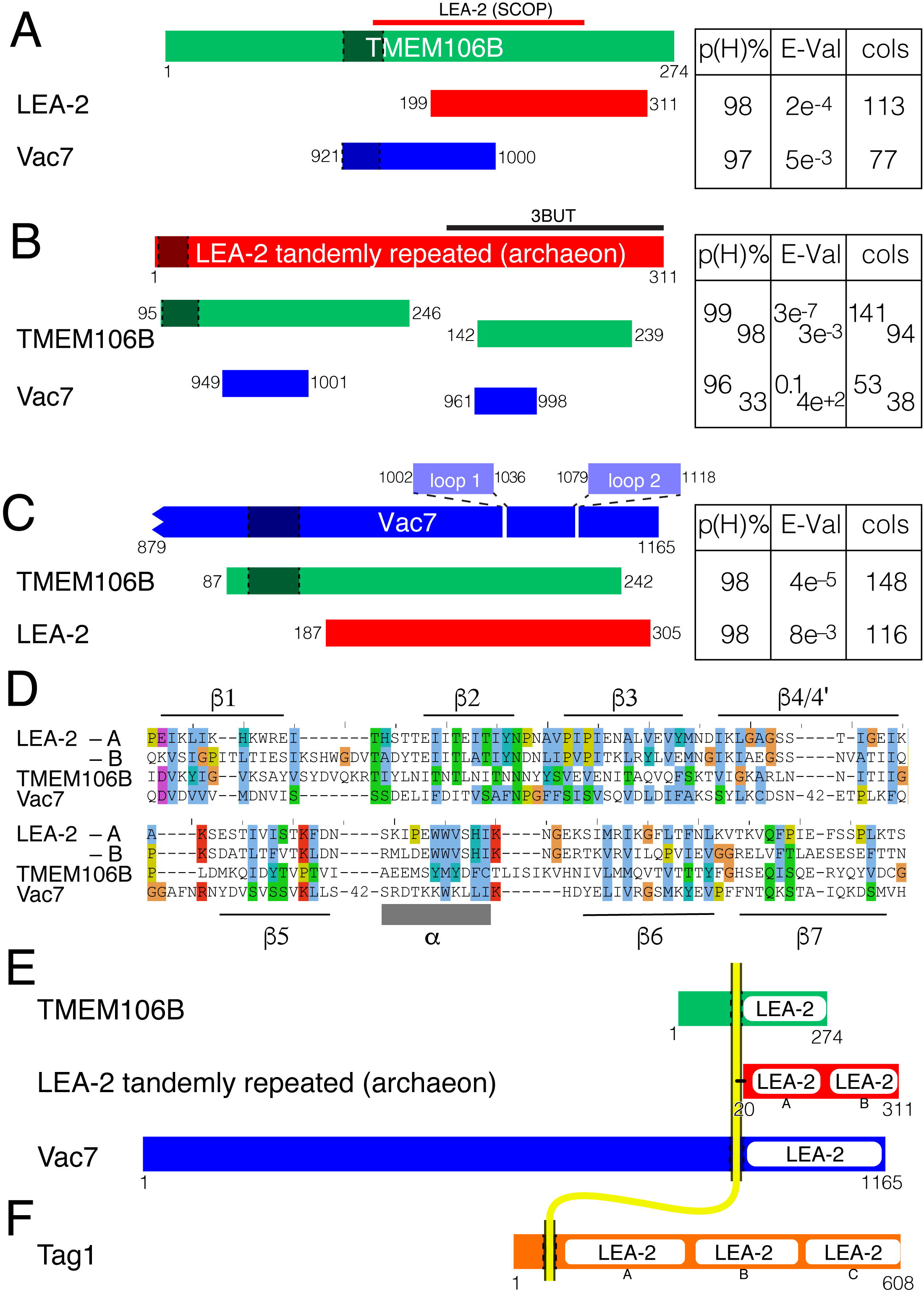
Homology between TMEM106B, LEA-2 and Vac7 identified by HHpred. A-C: Top hits from seeding HHpred with A. human TMEM016B (green); B. archaeal LEA-2 protein (*T. litoralis* WP_148290494.1, red, with tandemly repeated); C. the C-terminus of budding yeast Vac7 (blue). The regions of top hits that aligned with these seeds are shown, with statistics of the probability of homology p(H)%, the expected value that chance hits with a score better than this would occur if the database contained only hits unrelated to the query (E-Val), and the number of columns matched (cols). For TMEM106B as seed (A), the region of homology identified by SCOP is indicated above. For LEA-2 as hit (A and C) although HHpred made the hit to the solved structure of an *Archaeoglobus fulgidus* LEA-2 fragment (PDB: 3BUT), numbering is for the full length *T. litoralis* protein. For Vac7 as seed (C), the unstructured N-terminus and two non-conserved loops (residues 1002-1036 and 1079-1118) were omitted (see Supplementary Figure 1). For all parts, MSAs made by PSI-BLAST, rather than standard HHblits, produced similar identifications of homology, though with marginally lower probabilities (not shown). D: Alignment of sequences of LEA-2 domains in archaeal LEA-2 domain A (41-163) and domain B (174-307), TMEM106B (121254), and Vac7 (948-1162 missing two 42 residue inserts). Colouring according to Clustal scheme, and showing secondary structural elements. Strand 4’ consists of 3 residues that form an extension of strand 4. E: Domain maps of mature TMEM106B, archaeal LEA-2 and Vac7, including their relationship to the membrane (yellow). In all parts, hydrophobic segments are indicated by darker regions between dashed lines. This region is predicted to be cleaved in mature LEA-2, with acylation of a cysteine of position 20.^30^ F: Domain map of Tag1.

In both searches the next strongest hit was the C-terminus of Vac7. Probabilities of homology were 97/96%, with E-values for the alignment based on sequence comparison alone of 5×10^-3^/0.1 (Figure 2A/B). For TMEM106B, the region of homology extended into the TMH. The hits to Vac7 were shorter than the full length hits between TMEM106B and LEA-2, aligning only with ~60 residues after the TMH (as far as residue 1000). To investigate this, we submitted the C-terminus of Vac7 to HHpred, including 40 aa of the cytoplasmic domain, the single TMH and entire lumenal domain. This produced strong hits (pHom = 98/96%) to TMEM106B and LEA-2 proteins, but in both cases the homology again only included ~60 residues after the TMH (Supplementary Figure 1A). A possible reason for this was found in the alignment of Vac7 with itself, which predicted seven β strands and one α helix, as found in LEA-2, but also two unstructured regions, the first starting at residue 1002 and the second, which is repetitiously anionic, starting at residue 1079. These inserts are specific to budding yeast as they are not represented in the consensus sequence (Supplementary File 2). To test if the non-conserved inserts prevented two regions of homology being joined together, we carried out HHpred searches with the yeast Vac7 sequence missing either loop. This lengthened the alignment with archaeal LEA-2 to the end of Vac7, but did not alter the alignment with TMEM106B (Supplementary Figure 1B/C). Omitting both Vac7 loops produced full-length hits to both TMEM106B and LEA-2 proteins, with probabilities that they are homologous at 98% and E-values for the alignment based on sequence comparison alone of 4×10^-5^/8×10^-3^ (Figure 2C). The loop-less Vac7 sequence also increased the number of LEA-2 hits in Jackhmmer than the wild-type sequence (not shown).

The HHpred hits were not only strong, they had no features associated with false positives,^35^ namely they did not arise in repetitive, low complexity regions, they were equally strong in either direction, they produced low E-values based on sequence alone, and they had the same structural elements: seven β-strands and a single helix after strand 5 (Figure 2D, Supplementary Files 1-3). Finally, these regions all shared a conserved motif: N-p-N (where the preference for proline in position 2 is partial) located after strand 2 (Supplementary Files 1-3), which likely constitutes an Asx tight turn.^53^. Other tools were used to confirm HMMER and HHsearch. All of FFAS, SWISS-MODEL and PHYRE2 made the same assignment that TMEM106B and Vac7 are members of the LEA-2 superfamily (not shown).^36,37,54^ This is strong evidence that TMEM106B and Vac7 are members of the LEA-2 superfamily, with the C-terminal intra-luminal domains of both these proteins consisting of LEA-2 domains (Figure 2E), with the same topology as the archaeal protein, although this is predicted by the Signal 5.0 tool to be anchored to the membrane by acylation of a conserved cysteine just after the hydrophobic region.^30^

### Other LEA-2 proteins include the yeast vacuolar protein Tag1

Alongside hits to domains documented as TMEM106B, LEA-2 or Vac7, both HMMER and HHpred searches identified other hits in two categories: (1) regions designated as belonging to another protein family (n=1300 in HMMER); (2) regions without any prior designation (n=3000 in HMMER) (Supplementary Table 3A). In the first category, the dominant domain was DUF3712 (132 residues), almost all of which are in fungi. DUF3712 proteins containing one copy are ~240 aa in length, but ~50% are longer than 440 aa, reaching to over 4000 aa, many of which contain multiple copies. Homology of DUF3712 with LEA-2 was supported by finding seven β-strands and a single helix in DUF3712, however the two domains are out of register, with DUF3712 starting at strand 4 of one LEA-2 and ending at strand 3 of the next (not shown). Looking at one of the longer proteins: UM15053 in the fungus *Ustilago maydis* has 15 LEA-2 domains repeated gaplessly that have an out-of-phase relationship with all six annotated DUF3712 domains (Supplementary Figure 1D). The presence of partial DUF3712 domains at the end of a run of LEA-2s (Supplementary Figure 1E) confirms that the boundaries defined for DUF3712 are most likely an annotation artefact rather than a genuine circular permutation.

Many of the proteins in category (2) above (without any annotated domains) showed homology to DUF3712, in that the majority of hits from the first round of a Jackhmmer search were DUF3712 proteins. An example of this is the *S. cerevisiae* protein Tag1, a type II integral membrane protein of the yeast vacuole, named for its role in <u>T</u>ermination of <u>A</u>utopha<u>G</u>y (Supplementary Table 3C).^55^ HMMER and HHpred searches with Tag1 indicated it consists of a concatemer of three regions homologous to DUF3712 (each overlapping with LEA-2), with strongest homology for the DUF3712 hit nearest to the N-terminus (E-value below 10^-8^ for residues 181-334) (Supplementary Figure 1E). Although the links to each of the three homology on their own were weaker than found for Vac7, searches for tandemly repeated LEA-2 domains, for example when HHpred was seeded with the archaeal LEA_2 protein from Figure 2, produced a stronger hit to Tag1 than to any other yeast or human protein, with probability of homology >99% across 245 residues and E-value 2×10^-8^. Other tools confirmed aspects of this finding about Tag1: FFAS identified Tag1 as a homologue of DUF3712, Phyre2 determined that both DUF3712 and the final repeat in Tag 1 (domain C) were closer to LEA-2 domains than to any other structure, and SWISS-MODEL modelled DUF3712 as being like LEA-2. Together with its predicted short unstructured N-terminal cytoplasmic domain and single TMH, these results indicate that Tag1 is a second LEA-2 protein in budding yeast (Figure 2F).

To examine whether Tag1 is more related to Vac7 than other LEA-2 superfamily members we created an inter-relatedness map for ~2000 members of the LEA-2 superfamily, clustered according to all-vs-all BLAST using the CLANS tool.^34^ The map showed that the plant bacterial/archaeal and fungal LEA-2 proteins form a core of greatest connectivity, with all of TMEM106B, Vac7 and Tag1/DUF3712 being less connected (Supplementary Figure 2). Of these, TMEM106B is the most connected, particularly to the fungal LEA-2 group. Vac7 has connections to all three core groups but at a lower level than TMEM106B, and there is one direct connection between Vac7 and TMEM106B. The main group of DUF3712 and Tag1 proteins show are connect to the core similarly to Vac7, however the budding yeast Tag1 protein is quite peripheral, and only indirectly connected to any other groups. Thus, clustering indicates that the Vac7 and Tag1 are not paralogues and that they have independent origins in the LEA-2 superfamily. The close relationship of TMEM106 to fungal LEA_2 proteins (Supplementary Table 2B) may derive from a relatively recent common ancestry.

### Independent structural prediction that TMEM106B, Vac7 and Tag1 have lumenal LEA-2 domains

To seek further confirmation that TMEM106B, Vac7 and Tag1 are homologous to LEA-2, structures were predicted with trRosetta, which determines which pairs of residues have co-evolved, then estimates the proximity of the side-chains of each pair, and finally uses proximity to fold proteins *ab initio*.^38^ Using artificial intelligence approaches, the trRosetta tool was the best performing publicly available structure prediction tool in the 2020 Critical Assessment of protein Structure Prediction-14 (CASP14) exercise.^56^

Here, trRosetta was set to ignore all solved structures. This is significant because there are three solved structures of LEA-2 domains in the Protein Data Bank (PDB). All show seven-stranded β-sandwiches, with the same overall form as the immunoglobulin fold, capped by a single helix between strands 5 and 6. This structure is highly conserved between archaea (PDB: 3BUT) and *Arabidopsis* LEA-2 (two close homologues, PDB: 1xO8 and 1YYC), even though there is only 14% sequence identity (Figure 3A).^52,57,58^

**Figure 3:**
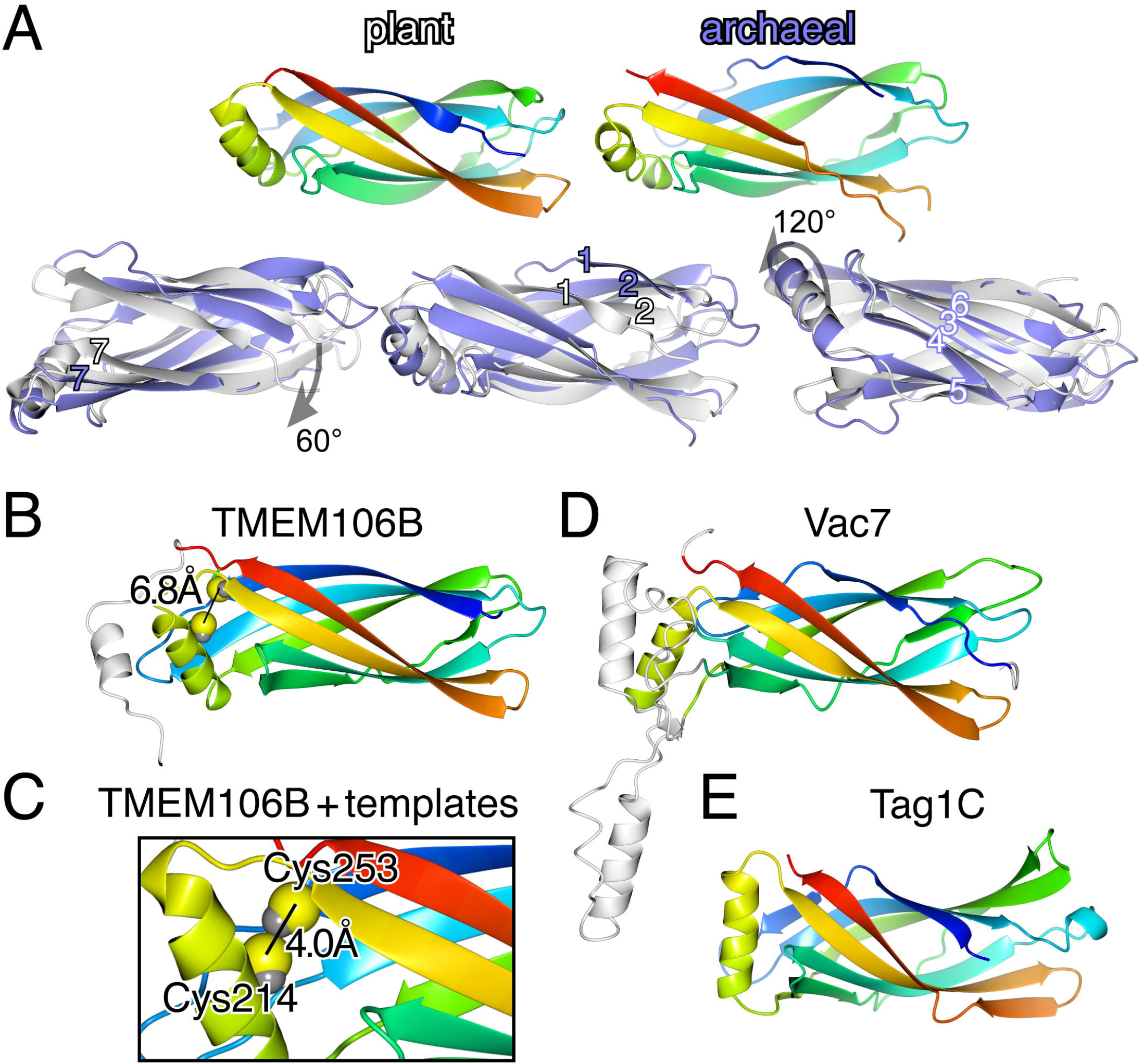
Structures of LEA-2 domains and predictions for the lumenal domains of TMEM106B and Vac7. A. 3D alignment of two known LEA-2 structures: plant (1XO8_A, residues 20-144, average of 20 NMR models) and archaeal (3BUT_A, residues 0-122, crystal). Top row: each LEA-2 structure shown in rainbow from N-to C-terminus. Bottom row: superposition of plant (white) and archaeal (light blue) in three views (centre: identical to top row; left: rolled 60° forward; right: rolled 120° backward). The seven strands are identified: for well aligned strands 3 to 6 - single numbers, white, blue surrounds); for less well aligned strands 1,2 and 7 - by white or blue numbers, both with black surrounds. Fog indicates increasing depth. B. Structure of the C-terminus of TMEM106B predicted by trRosetta without using solved structures as templates. Orientation and colouring as in A (top), with helix-2 at extreme C-terminus in white. Sulfhydryls of cysteines 214 and 253 are shown with the interatomic distance. C. Detail from structural prediction of TMEM106B that did use solved structures, with the interatomic distance of sulfhydryls C214 and C253. D. Predicted structure of the C-terminus of Vac7, ignoring solved structures as part B. Two elements do not align with LEA-2 (white): (i) loop between β3-4: includes a short predicted helix in a similar position to the additional one in TMEM106B; (ii) extended loop between β5-helix: includes a short helix, position variable. E. Predicted structure of C-terminal region of Tag1 (domain C), coloured as part B.

For TMEM106B, trRosetta predicted a LEA-2-domain with very high confidence (Supplementary Table 4). Pairwise amino acid co-evolution identified the major regions of contact as five pairs of anti-parallel β-strands (Supplementary Figure 3). The TMEM106B model aligned closely with archaeal LEA-2, with an additional helix at the extreme C-terminus, the orientation of which was uncertain (varied in top 5 models, not shown) (Figure 3B). TMEM106B has two conserved cysteines (C214 and C253, Supplementary File 1A), and the model placed them close together (Figure 3B). In a further trRosetta model that included solved structures as templates, both cysteines shifted slightly making the sulfhydryls touch (inter-atomic distance 4.0Å, Figure 3C). This strongly suggests that the TMEM106B structure is maintained by a conserved disulphide bond. This may be clinically relevant, as mutation in an adjacent residue (D252N) causes hypomyelinating leukodystrophy type 16, paralleling the phenotype of complete gene loss.^7,8^

For Vac7, the model was predicted with high confidence (Table 3), or with very high confidence if the two loops were emitted (not shown). The model aligned closely with LEA-2, although the helix between strands 5/6 was predicted to lie in an orientation 45° different from that of archaeal LEA-2 and TMEM106B. The two inserts in Vac7 are between strands 3 and 4 (42 residues) and between strand 5 and the helix (38 residues) (Figure 3D). The modelled position of the first insert was similar to the additional helix in TMEM106B, while the position of the second insert was variable (not shown).

For Tag1, domain C was strongly predicted as like LEA-2 (all 5 top models) (Figure 3E), while domain B was like LEA-2 only in three models, and domain A was a β-sandwich, but did not contain strands 6 and 7 in any model (not shown). There were also minor variations among the predicted LEA-2-like domains, with extra helices in domain A, and strands 1 and 7 in domain B replaced by short helices (not shown).

For predicted structures of all five newly predicted LEA-2 domains (TMEM106B, Vac7 and the three domains in Tag1) the closest hit among solved structures in PDB (nr25 subset) was the archaeal LEA-2 crystal structure 3BUT.^39^ A matrix of pairwise comparisons between modelled LEA-2 domains showed that TMEM106B and Vac7 were more similar to each other than either was to domain C of Tag1 (Supplementary Table 4).

### LEA-2 is a lipid transfer protein

Although the archaeal LEA-2 structure (3BUT) was released in the protein database (PDB) in 2008, it has never been described in any report.^52^ Inspection shows that its β-sandwich splays apart to create a groove between strands 1 and 7, dimensions 27Å long, 6Å wide and 7Å deep that is largely hydrophobic and highly conserved (Figure 4A/B). In the crystal, the groove contains an almost unbroken chain of 10 water molecules (Figure 4C). This was observed previously in the PDB file for 3BUT, which contains this remark: “[an] undefined ligand or cofactor is bound into the central cavity, a part of it is most likely a lipid. This ligand has not been modeled”.^52^ One end of the groove is closed off, with its base formed by conserved side chain γ methyls of the final residue of strand 4 (V59, Supplementary Figure 4A/B). At the other end, beyond the chain of waters, the groove widens out to a conserved hydrophobic indentation ~10Å in diameter (Figure 4A/B, Supplementary Figure 4C/D).

**Figure 4:**
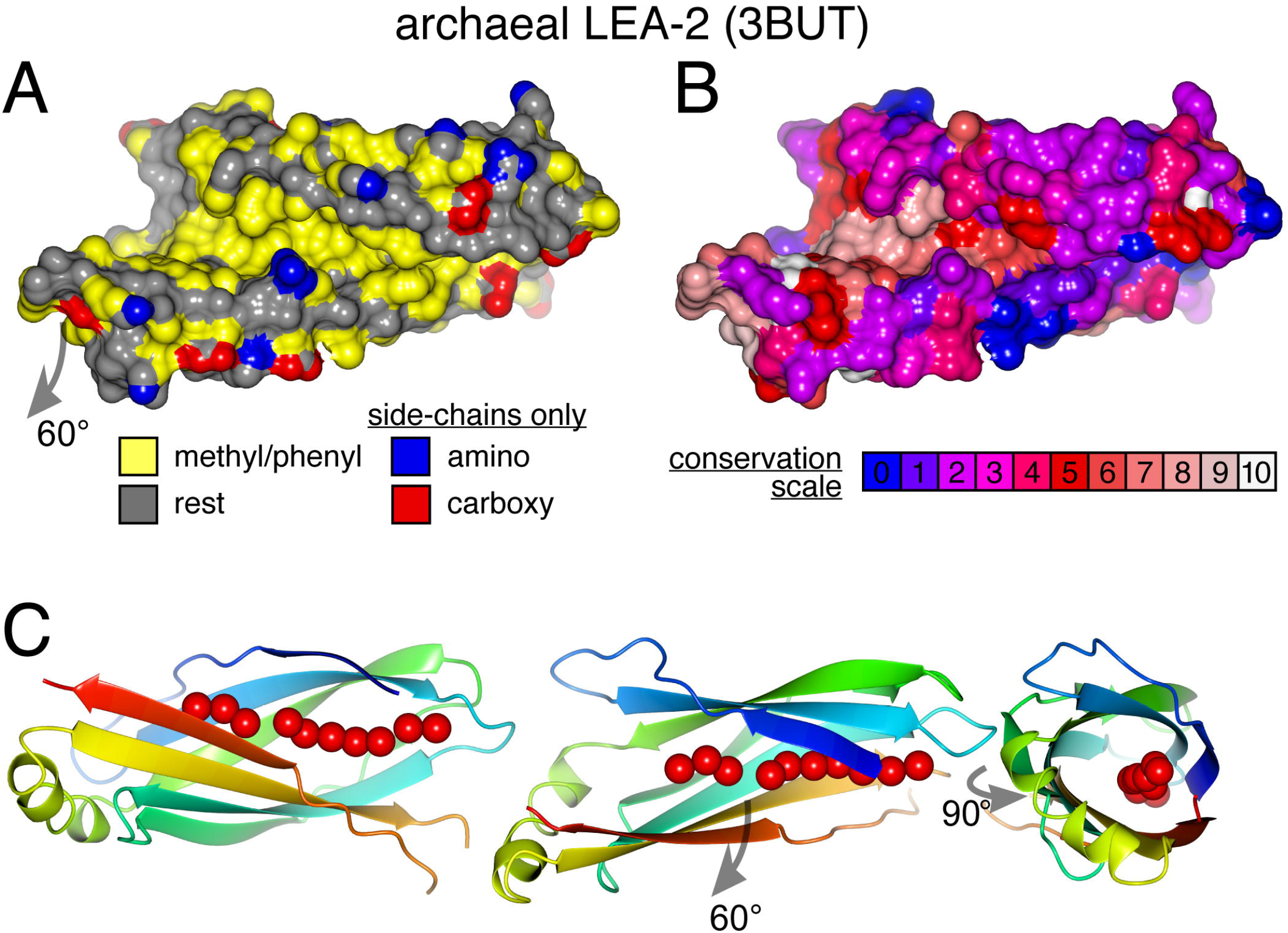
LEA-2 domains have a conserved lipid binding hydrophobic groove. A. Archaeal LEA-2 (3BUT) with surface coloured to highlight hydrophobicity and charge using the YRB scheme, with key.^40^ Position same as Figure 3A - rolled forward 60°. B. Archaeal LEA-2 (3BUT) with surface colour indicating conservation, according to the key (see Methods). C. Archaeal LEA-2 (3BUT) showing 10 water molecules (red spheres) in a longitudinal groove between strands 1 and 7. Protein colouring as Figure 3A. Centre: position as in A, with additional views: left - rolled back 60° into initial position from Figure 2; right - open end of groove side swung forward 90°.

The finding of a groove is not universal in LEA-2 structures, as it is not seen in the NMR structures of plant LEA-2 proteins (1XO8 and 1YYC), even though the residues that line the groove are conserved (not shown). Looking inside the plant LEA-2 structures, both contain a series of internal cavities running down the centre of the domain towards the conserved hydrophobic residue (Supplementary Figure 5A/B). Among the models of newly predicted LEA-2 domains, neither of the TMEM106B and Vac7 models had a groove, but both contained cavities like 1XO8 and 1YYC (Supplementary Figure 5C/D). Only domain C from Tag1 contained a surface groove (Supplementary Figure 4E). As a control, models of other immunoglobulin fold β-sandwiches, for example an Ig light chain constant region, did not contain a string of cavities (Supplementary Figure 5E).

Overall, the hydrophobic surface and dimensions of the groove in 3BUT suggest that LEA-2 domains solubilise an extended hydrophobic molecule such as a fatty acid, a lysolipid or possibly a diacyl bilayer lipid. If this structural property is verified, the LEA-2 superfamily, including TMEM106B Vac7 and Tag1, would be classified as lipid transfer proteins.^59,60^ The variability in finding a groove in different structures is addressed in the Discussion.

### The TMHs of TMEM106B and Vac7 are homologous while the cytoplasmic domains are divergent

While the lumenal domains of TMEM106B and Vac7 are homologous, other portions of the proteins may have evolved differently, which would lead the homologues to adopt distinct functions. This makes it worthwhile to survey the sequence features of the other regions.

The cytoplasmic domains of TMEM106B Vac7 and Tag1 are predicted to be almost entirely unstructured, even though for Vac7 this is >900 residues (not shown).^20,50^ MEME detected multiple conserved motifs in the N-terminus of Vac7,^61^ but none are shared with TMEM106B or Tag1 (not shown). The N-terminus of TMEM106B (and TMEM106A/C) starts with a predicted myristylation signal (MGxxxS), which would promote membrane anchoring.^62^ TMEM106B also contains the motif CxxCxGxG. Since two of these can form a zinc-binding site, this provides a second means by which TMEM106B can dimerise.^63^ Similar cysteine-rich motifs are found once or twice in the N-termini of many plant and fungal LEA-2 proteins, but not in the families of Vac7, Tag1 or archaeal LEA-2.

Considering just the TMHs, their sequence properties are conserved across evolution within and between the families. The similarity is such that HHpred search seeded with 40 residues from the TMH of either TMEM106B, Vac7 or fungal LEA-2 produced hits to plant LEA-2s (not archaeal) above all non-self proteins in humans yeast and *Arabidopsis*, while this was not the case for the TMH of Tag1 (not shown). The similarities within TMHs arose from two conserved features: (i) a cluster of positive residues mixed with small residues at the cytoplasmic end (RLRPRRTK for TMEM106B, NINNRHKK in plant LEA-2 At5g53730, RKSPFVKVKN in Vac7); and (ii) dimerising σxxxσ motifs, where σ is a residue with a small side-chain (G, A, S or T).^64^ The TMH of TMEM106B has S/AxxxCxxxSG/S, reminiscent of dimerising motifs of the form SxxxCS in Plexin-D1, Vac7 has GxxxG, and similar motifs are found in plant and fungal LEA-2 proteins (for example: *Arabidopsis* At5g53730 has STxxSG, *Rhizophagus* A0A2H5R616 has GxxxA).^65^ Such motifs are absent from tag1 and its closest homologues (not shown). Thus, while the cytoplasmic domains are divergent in length and in sequence, the TMHs of TMEM106B, Vac7 and eukaryotic LEA-2 proteins promote dimerisation.

## Discussion

The predictions of homology between the C-terminal domains of TMEM106B and LEA-2, and then between LEA-2 and two yeast proteins, Vac7 and Tag1, arose by applying different sequence comparison tools. The link between TMEM106B and LEA-2 is so solid it can be made with PSI-BLAST. The link to Vac7 required more sensitive tools (HMMER, HHpred). Knowing the LEA-2-Vac7 link, the PSI-BLAST searches seeded with Vac7 were reexamined and found to contain a small number of LEA-2 proteins, although most were below its significance threshold (Supplementary Table 5). Although LEA-2-Vac7 homology may be too distant to be detected by low sensitivity tools, it is supported by many approaches. The same applies for Tag1, particular when searches were seeded with tandemly repeated LEA-2 domains.

The predicted fold for TMEM106B, Vac7 and all three domains of Tag1 was corroborated by the independent approach of contact folding using trRosetta. Even set to ignore known structures as templates, the closest solved structure for all these domains was a LEA-2 domain (not shown). In addition to the similarity between these domains, the topology of the proteins and their intracellular localisations to late endosomes and lysosomes/vacuoles are similar, indicating a shared origin and some aspects of shared function at the molecular level.

Among these three new LEA-2 proteins, TMEM106B and Vac7 share the phenotype of lysosome/vacuole enlargement, however this may work in opposite directions. For Vac7, vacuolar enlargement accompanies deletion.^50,66^ In contrast, lysosomal enlargement is associated with over-expression of TMEM106B, and deletion has no effect on the bulk of lysosomes.^15–18^ The conflicting phenotypes raise the possibility that TMEM106B and Vac7 have evolved in different directions from a common ancestor.

At the molecular level, more is known about Vac7 than TMEM106B. Vac7 is required for stress responses that increase the late endosomal/ lysosomal inositide lipid PI(3,5)P_2_.^50^ It is still not established if Vac7 achieves this by activating the PI3P 5-kinase Fab1 that synthesises PI(3,5)P_2_, or by inhibiting Fig4, the PI(3,5)P_2_-phosphatase. These opposing lipid-modifying enzymes are members of a single complex, scaffolded by Vac14.^67–69^ The tripartite complex is conserved widely in eukaryotes, including humans, where the kinase is called PIKfyve and the other two proteins retain their yeast names. The mechanism of action of Vac7 does not involve altering the assembly or membrane targeting of the Fab1 (PIKfyve) complex with Vac14 and Fig4.^67,70^ Nevertheless, the cytoplasmic domain of Vac7 strongly interacts with Vac14.^72^ The interaction interface requires almost the whole of Vac14, which has different binding sites along its length,^69^ which suggests that Vac7 may bind not to Vac14 alone, but also to partners of Vac14.

Turning to TMEM106B, while its over-expression or deletion causes wide-ranging effects on lysosomes,^15–19^ it is not known which of these are primary. The breadth of effects is consistent with it affecting production of PI(3,5)P_2_, which recruits many effectors as is common for most inositide lipids.^75,76^ Other evidence links TMEM106B to PI(3,5)P_2_ levels: TMEM106B is a top hit for host proteins required for SARS-CoV2 infection,^12–14^ which also requires PIKfyve and Vac14.^13,14,77^ Finally, in *Trichinella* nematode worms the open reading frames for TMEM106B and Fig4 are positioned so close to each other that they are annotated as a single TMEM106-Fig4 fusion protein. This appears likely to be an error, as it places Fig4 in the lumen (not shown). More likely, TMEM106B and Fig4 are co-regulated in one of the many bicistronic operons in this species.^78^ Such co-regulation suggests that Fig4 might be a binding partner for the N-terminus of TMEM106B.

Turning to possible molecular functions for the new LEA-2 proteins, the archaeal crystal structure has an obvious lipid binding groove. Although the groove is missing from two NMR structures of plant LEA-2 proteins, the residues required to form the groove are conserved across the whole superfamily, and the NMR structures and every LEA-2 domain that could be modelled in its entirety contain an array of internal cavities along the same line as the groove (Supplementary Figure 5). These cavities were not reported in the single paper about these structures,^57^ and their significance is unknown, but one hypothesis is that they indicate an “apo” (empty) conformation of LEA-2 domains, while 3BUT is closer to a “holo” (ligand bound) conformation, consistent with the finding that the crystal contained an undefined, unmodelled ligand.^52^ This would imply that LEA-2 domains undergo a conformational change that parallels other lipid transfer proteins, where conformational change either allows lipid entry into a deep pocket,^79,80^ or is necessary to accommodate lipid.^81–83^ Given the findings that LEA-2 domains have the features of lipid transfer proteins, the predictions that TMEM106B, Vac7 and Tag1 have lumenal LEA-2 domains links them to lipid metabolism in an as yet unknown way. Based on analogy with other lipid transfer proteins, there are three possible modes of action downstream of lipid solvation: presenting lipid from the membrane to a lumenal enzyme (similar to lysosomal saposins); transferring lipid from intra-lysosomal vesicles or lipoproteins to the limiting membrane (similar to lysosomal Niemann-Pick type C protein 2); or sensing lipid by changing intra- or inter-molecular interactions in response to lipid binding (like nuclear StART domains).^60,84^ Tag1 may be informative on the mode of action of TMEM106B or Vac7, even though it is the most variant of the new LEA-2 proteins (Supplementary Figure 2). In the sole report on Tag1, it was found to respond to prolonged starvation by migrating to a small number of spots in the vacuolar membrane, from where it signals to inhibit cytosolic Atg1, the yeast homologue of ULK1, thus terminating autophagy.^55^ Accumulation in spots and signaling function required the entire lumenal domain and membrane attachment, which could not be reconstituted with non-Tag1 elements. This suggested a model that Tag1 senses a signal derived from autophagic material that builds up during starvation and communicates the signal to terminate autophagy. In the model, amino acids were proposed as a plausible homeostatic signal.^55^ Speculatively, might it be that the signal instead is lipid-based, and also that both Vac7 and TMEM106B (and maybe many more LEA-2 proteins) respond like Tag1 to lipid signals and transmit them to partners in the membrane or on the cytosolic side?

Overall, this study reveals homology of TMEM106B in animals with Vac7 and Tag1 in yeast, and suggests unanticipated molecular behaviour that they might share. However, the study is limited in that it says nothing about which hydrophobic molecules bind the predicted LEA-2 domains or how this changes the behaviour of the full-length proteins. Despite these issues, modelling TMEM106B, Vac7 and Tag1 as lipid transfer proteins will guide future experiments that test the function of these proteins, for example through point mutations designed to inhibit lipid uptake, which might be achieved by filling the lipid binding groove with large hydrophobic side-chains.^85^

## Supporting information

All supplementary Material (5 Figs, 5 Tables)

## Data availability statement

The data that support this study are freely available in Harvard Dataverse at https://dataverse.harvard.edu/dataverse/LEA_2.

## Acknowledgements

work was funded by the Higher Education Funding Council for England and the NIHR Moorfields Biomedical Research Centre

