## Supplementary material for "TMEM106B in humans and Vac7 and Tag1 in yeast are predicted to be lipid transfer proteins": All supplementary Material (5 Figs, 5 Tables)

**(a)** Supplementary Figures 1-5: pages 1-7
**(b)** Supplementary Tables 1-5: pages 8-10

#### (a) Legends for Supplementary Figures

TMEM106B in humans and Vac7 and Tag1 in yeast are predicted to be LEA-2-like lipid transfer proteins in lysosomes/vacuoles

**Supplementary Figure 1: Non-conserved loops in Vac7 reduce the strength of hit between Vac7 and TMEM106B/LEA-2.**

A-C: Top hits from HHpred searches with Vac7 879-1165 (blue): A. complete, B. lacking loop 1002-1036, C. lacking loop 1079-1118. Hits show the aligned regions of human TMEM016B (green) and archaeal LEA-2 protein (3BUT structure, using homologous numbering for *T. litoralis* WP_148290494.1, red), with statistics of the probability of homology p(H)%, the expected value that chance hits with a score better than this would occur if the database contained only hits unrelated to the query (E-Val), and the number of columns matched (cols). D: Domains in *Ustilago maydis* protein UM15053 (2590 aa), with each domain predicted from HHpred searches. Pink rectangles indicate hydrophilic regions with predicted mixture of short helices and unstructured loops. E. Alignment of Tag1 with both archaeal LEA-2 (as in A to C) and the protein family DUF3712 (132 residues), with statistics of each hit as in A to C.

**Supplementary Figure 2: Cluster map for LEA-2 Superfamily**

1951 sequences related to TMEM106B, Vac7, DUF3712 and Tag1 were connected by all-vs-all BLAST, and connections were used to cluster proteins according to their similarity to others (see Methods). Clusters are colour coded: black = plant, yellow = prokaryote, green = fungal LEA-2, red = TMEM106B, blue = Vac7, cyan = DUF3712, purple = Tag1, grey =other. Darker lines imply BLAST hits of increasing significance. The position of Tag1 in budding yeast is indicated by a large filled circle, while TMEM106B and Vac7 are in the cores of their respective families (not indicated).

**Supplementary Figure 3: Metrics of pair-wise co-evolution for TMEM106B.**

A. Contact map, displaying the predicted probability (0 to 1 scale) that residue pairs are in contact, which is defined as a gap of 8Å or less between their C-beta (C-alpha for Glycine) atoms. Outlined squares on the long diagonal indicate nine secondary structural elements, which are listed across the top. Filled squares in the top right half indicate which anti-parallel β-strand pairs show close contact. B. Distance map, showing the predicted proximities between residue pairs up to 20Å as per the scale. Both diagrams were created by trRosetta for residues 118-274 of TMEM106B, without using solved structures as templates.

**Supplementary Figure 4: Closed off end of hydrophobic groove in Archaeal LEA-2**

Views of archaeal LEA-2 (3BUT) from along the groove, with the domain rotated 15° backward compared to Figure 3A (A and B), and rolled forward 60° with the open end of groove side swung forward 75° (C and D), with surface highlighting of hydrophobicity and charge (A/C) or conservation (B/D), with colouring as in Figure 4A and 4B respectively. Water molecules are shown as small black spheres. In A and B, the surface at the apex of the blocked off end of the groove is made of the two γ methyls of V59. E Views of the model of Domain C in Tag1, coloured as A and C, with hydrophobic groove indicated by open arrows.

**Supplementary Figure 5: Internal cavities in LEA-2 domain structures.**

10Å slices through super-immunoglobulin domains, with all surfaces, including internal cavities, coloured black, and the side-chains of the hydrophobic residue at the end of strand 4 coloured yellow. A. 1XO8 and B. 1YYC are both plant LEA-2 structures solved by NMR. C. and D. are models from trRosetta (without templates) of TMEM106B and Vac7. E is from an immunoglobulin light chain (1RZI_A, residues 110 -213, highlighting L175). Other, 4+3 β-sandwiches in the Ig superfamily without internal cavities included β-galactosidase (region near L283 in 6QUC_A) and the chaperone CupB2 (region near V206 in 3Q48_B).

### Supplmentary Figure 1

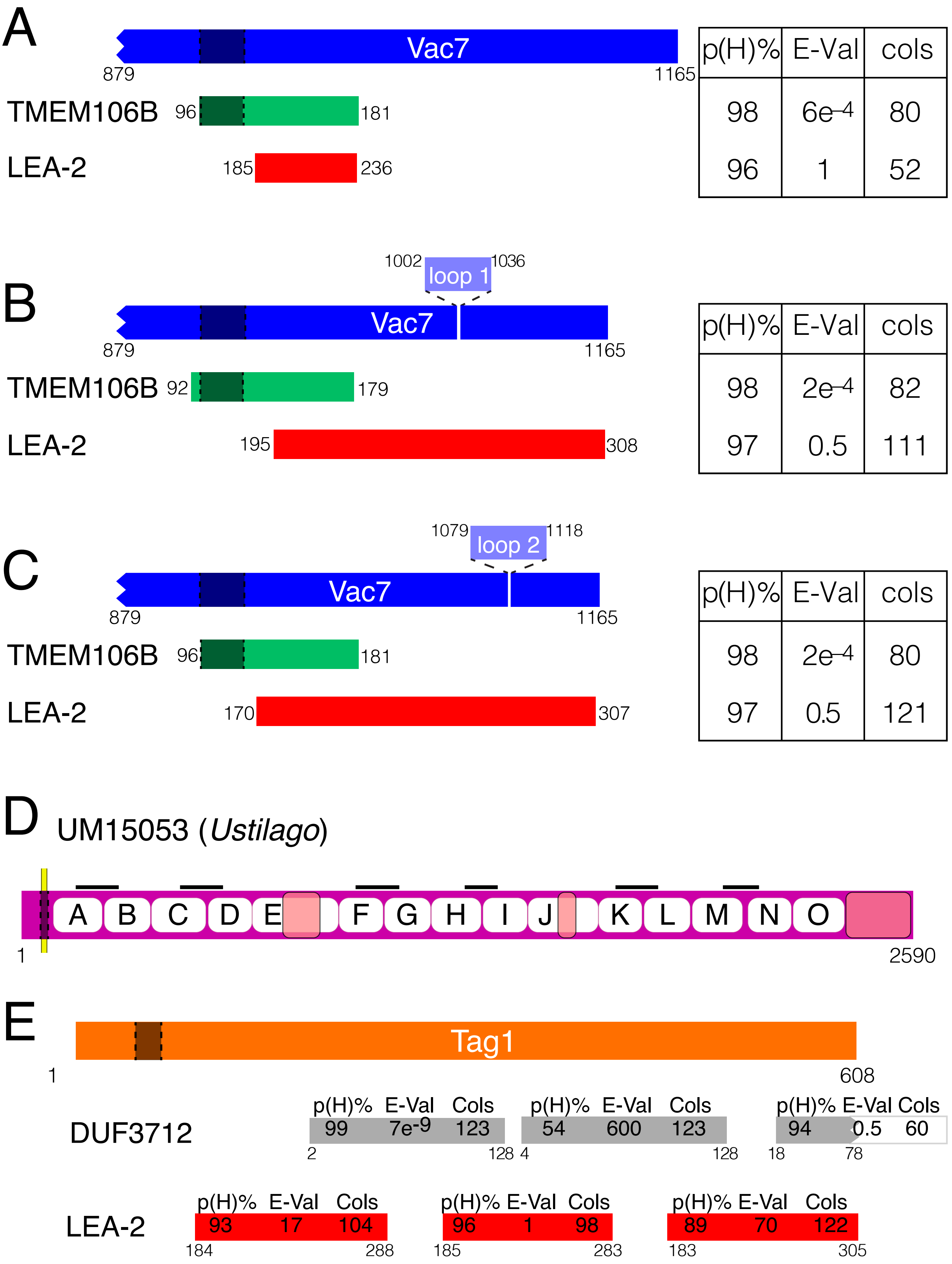

### Supplementary Figure 2

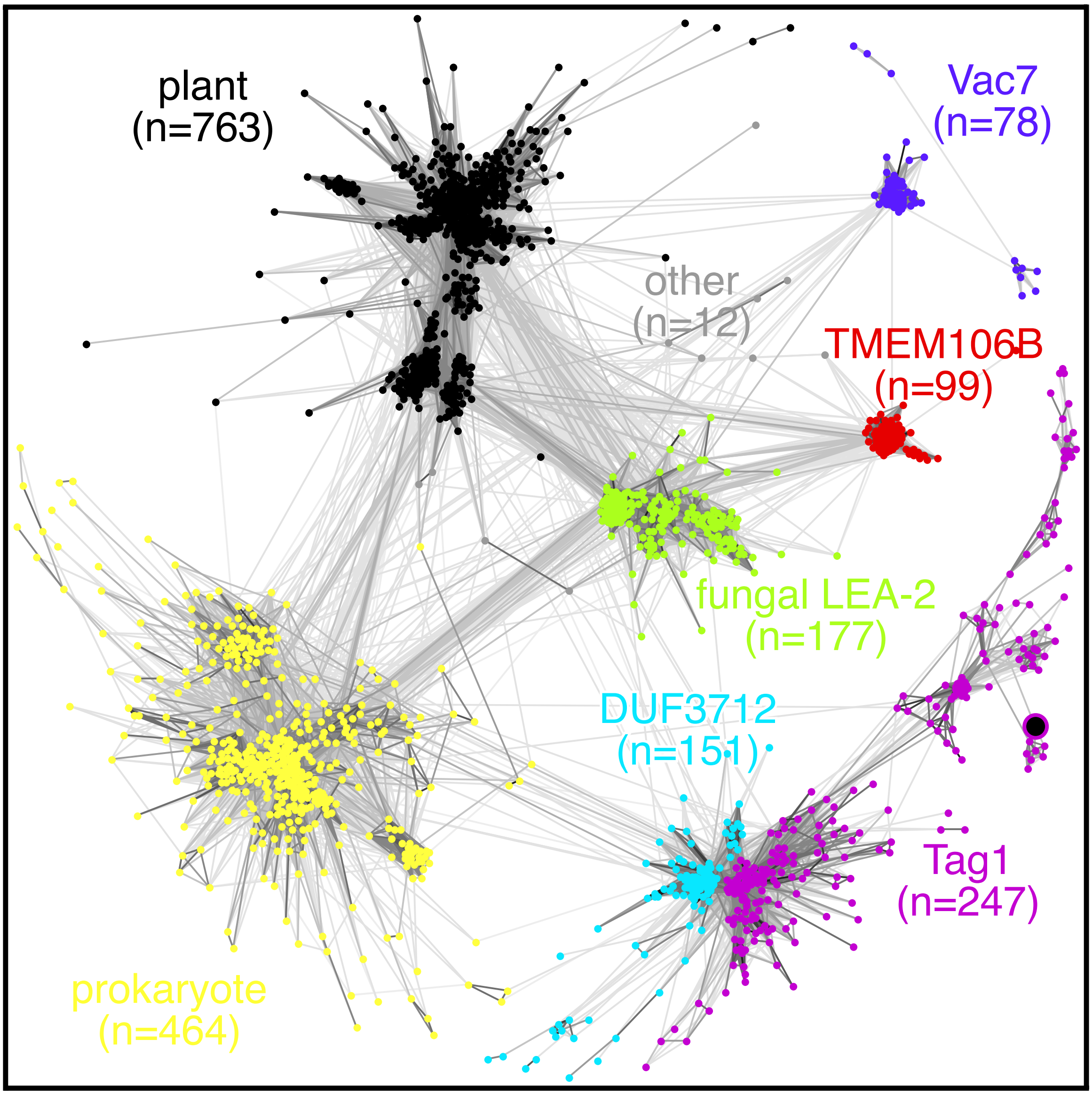

### Supplementary Figure 3

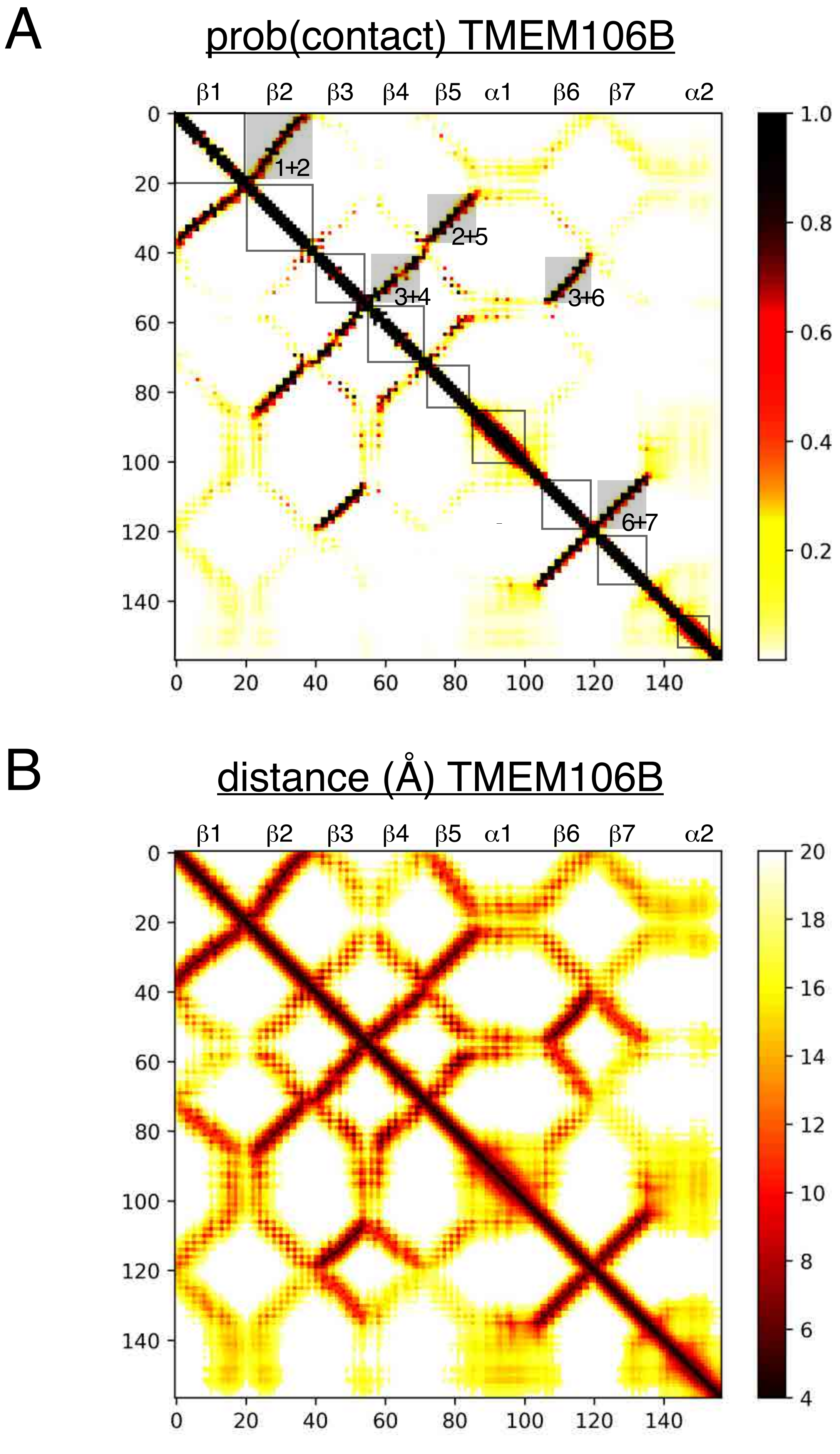

### Supplementary Figure 4

archaeal LEA-2 (3BUT)

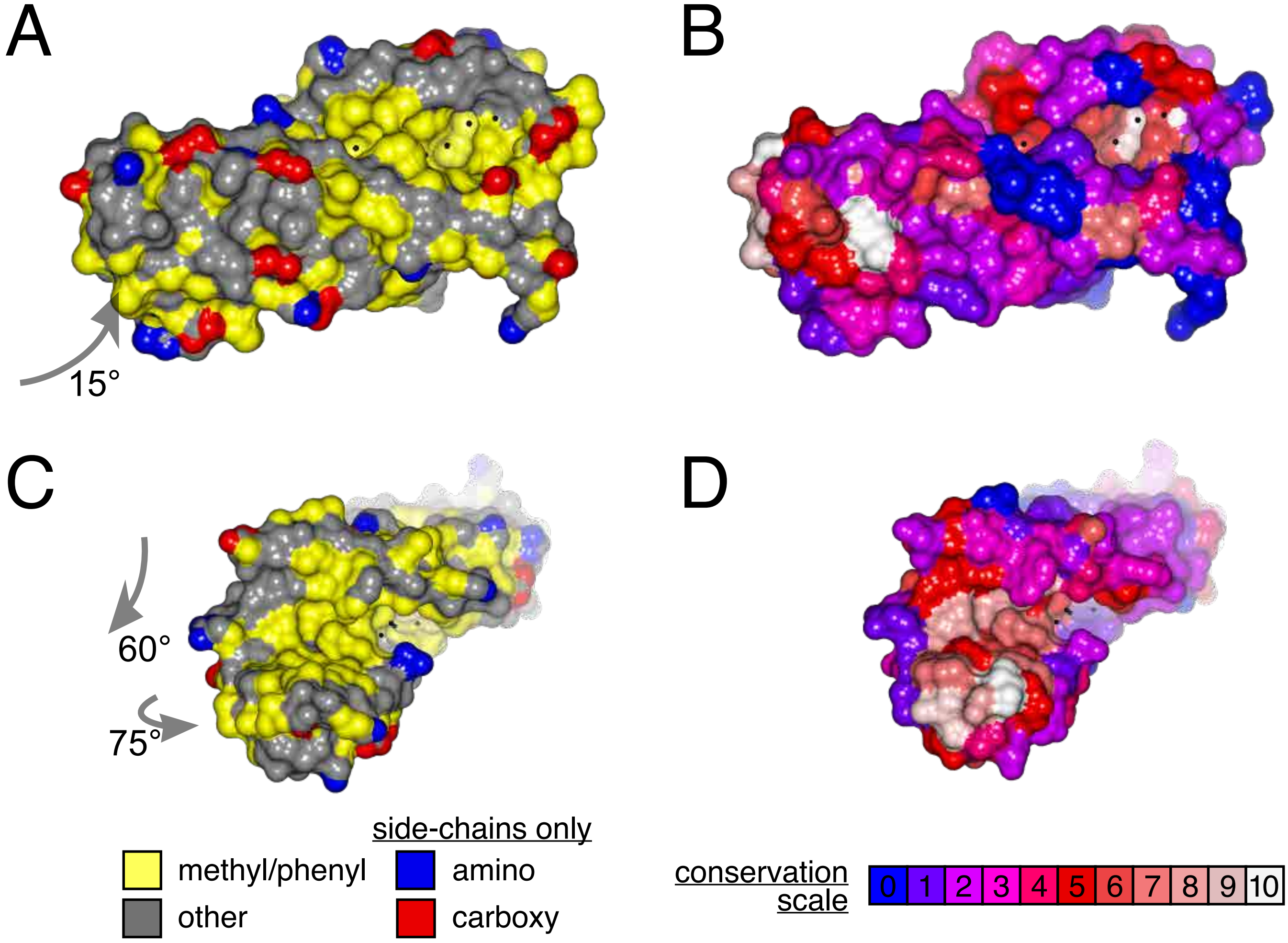


E

Tag1 domain C

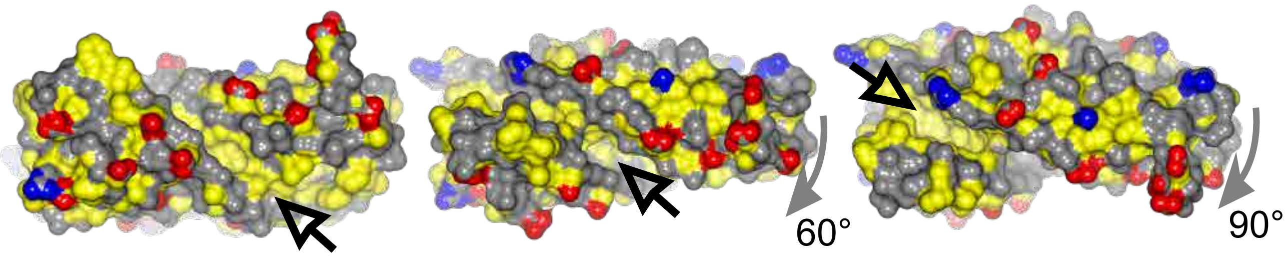

### Supplementary Figure 5

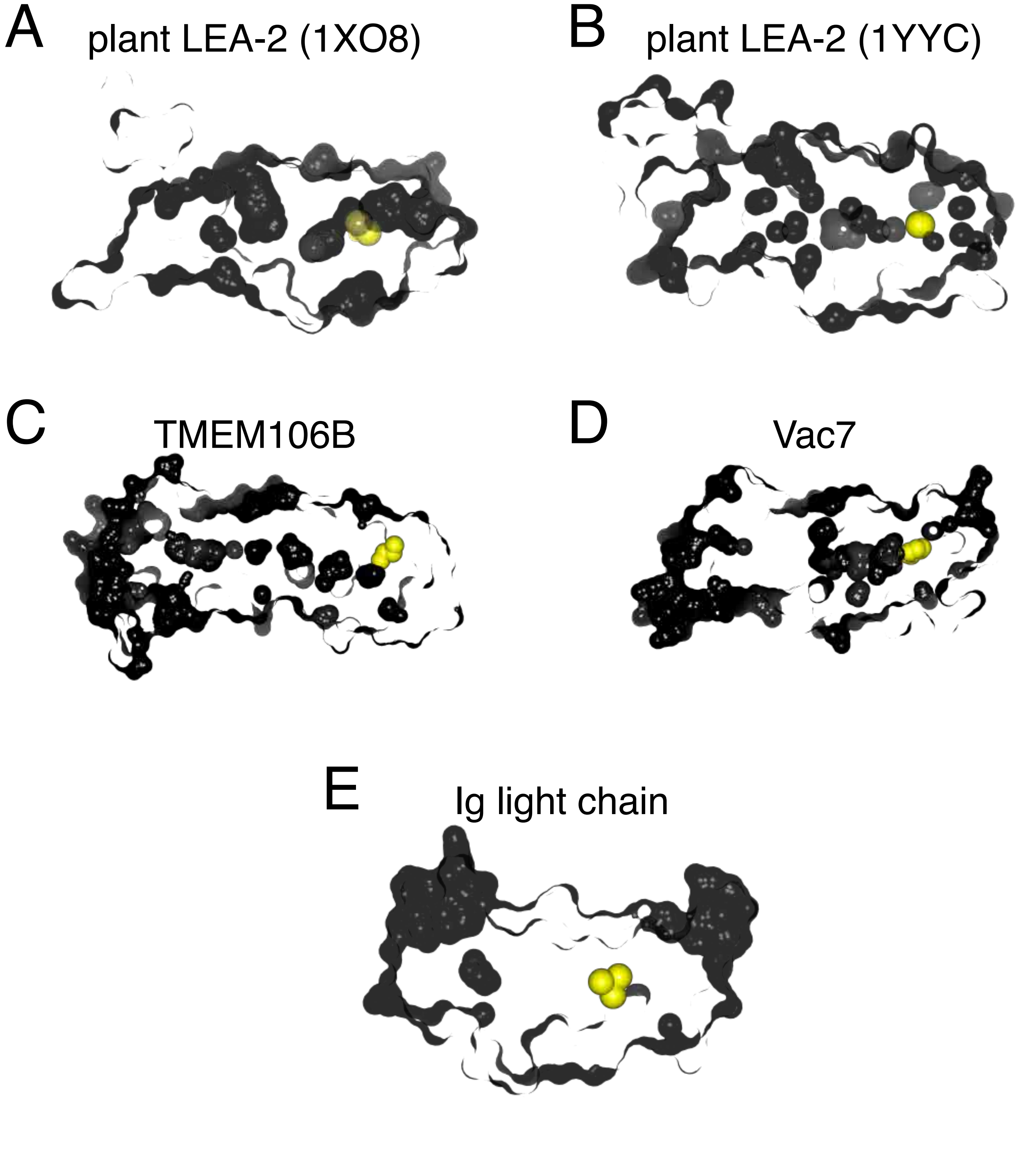

#### (b) Supplementary Tables

##### Supplementary Table 1. Two PSI-BLAST strategies to study TMEM106B

| database | iteration | total hits | TMEM106/ DUF1356 | | no domain identified | | | LEA-2 | |
|---|---|---|---|---|---|---|---|---|---|
| | | | only | with others | TMH | unknown | no info | named LEA-2 | other name |
| nr100 | 1 | 2486 | 2178 | 33 | 31 | 234 | 10 | 0 | 0 |
| | 2 | 2522 | 2181 | 35 | 31 | 258 | 17 | 0 | 0 |
| | 3 | 2523 | 2182 | 35 | 31 | 259 | 16 | 0 | 0 |
| nr50 | 1 | 173 | 121 | 3 | 5 | 28 | 16 | 0 | 0 |
| | 2 | 215 | 131 | 3 | 5 | 25 | 39 | 12 | 0 |
| | 3 | 381 | 135 | 3 | 5 | 66 | 44 | 127 | 1 |
| | 4 | 736 | 142 | 3 | 6 | 169 | 45 | 355 | 16 |
| | 5 | 844 | 109 | 2 | 4 | 212 | 12 | 477 | 28 |
| | 6 | 1975 | 149 | 3 | 7 | 600 | 26 | 978 | 212 |

Hits to TMEM106B identified using the two PSI-BLAST strategies were analysed for domain composition as described in Methods.

##### Supplementary Table 2: Domains in DUF1356, LEA-2, Vac7 and DUF3712 families across evolution.

A. Proteins with domains of interest in different taxa. NCBI database only.

| | DUF1356 | LEA-2 | Vac7 | DUF3712 |
|---|---|---|---|---|
| plants | 30 | 12557 | 149 | 1 |
| animals | 2540 | 16 | 203 | 2 |
| fungi | 18 | 436 | 1687 | 2854 |
| other eukaryotes | 6 | 42 | 47 | 370 |
| bacteria | 391 | 9224 | 192 | 11 |
| archaea | 1 | 830 | 21 | 3 |
| total | 2986 | 23014 | 2292 | 3241 |

B. Domains in fungi only, comparing PFAM, Uniprot and NCBI databases

| domain identified in fungi | database | total | dikarya | | zygomycete | | zoosporidia | | others |
|---|---|---|---|---|---|---|---|---|---|
| | | | asco-mycota | basidio-mycota | mucor-mycota | zoopago mycota | chytridio-mycota | blasto cladio-mycota | |
| TMEM106B (DUF1356) | PFAM | 0 | 0 | 0 | 0 | 0 | 0 | 0 | 0 |
| | Uniprot | 0 | 0 | 0 | 0 | 0 | 0 | 0 | 0 |
| | NCBI | 18 | 4 | 3 | 2 | 5 | 4 | 0 | 0 |
| LEA-2 | PFAM | 74 | 1 | 49 | 21 | 2 | 1 | 0 | 0 |
| | Uniprot | 109 | 2 | 75 | 28 | 2 | 2 | 0 | 0 |
| | NCBI | 436 | 46 | 222 | 142 | 13 | 10 | 3 | 0 |
| Vac7 | PFAM | 638 | 560 | 0 | 64 | 6 | 6 | 0 | 2 |
| | Uniprot | 910 | 817 | 0 | 72 | 9 | 10 | 0 | 3 |
| | NCBI | 1687 | 1513 | 83 | 72 | 8 | 8 | 0 | 3 |
| DUF3712 | PFAM | 1995 | 997 | 580 | 379 | 7 | 22 | 2 | 8 |

**Supplementary Table 3: Links between LEA-2 superfamily domains revealed by Jackhmmer**

**A. TMEM106B**

| iteration | Total | TMEM106B | LEA-2 | Vac7 | DUF3712 | other | none |
|---|---|---|---|---|---|---|---|
| 1 | 824 | 816 | 4 | 0 | 0 | 0 | 4 |
| 2 | 1163 | 837 | 279 | 0 | 0 | 0 | 47 |
| 3 | 4075 | 839 | 2620 | 0 | 34 | 21 | 561 |
| 4 | 9408 | 824 | 6237 | 15 | 366 | 74 | 1892 |
| 6 | 12382 | 824 | 6964 | 331 | 1042 | 245 | 2976 |

**B. Vac7**

| iteration | Total | TMEM106B | LEA-2 | Vac7 | DUF3712 | other | none |
|---|---|---|---|---|---|---|---|
| 1 | 519 | 0 | 0 | 515 | 0 | 0 | 4 |
| 2 | 537 | 0 | 1 | 528 | 0 | 0 | 8 |
| 3 | 548 | 0 | 12 | 528 | 0 | 2 | 38 |
| 4 | 939 | 0 | 371 | 528 | 0 | 2 | 38 |

**C. Tag1**

| iteration | Total | TMEM106B | LEA-2 | Vac7 | DUF3712 | other | none* |
|---|---|---|---|---|---|---|---|
| 1 | 227 | 0 | 0 | 0 | 127 | 0 | 100 |
| 2 | 629 | 0 | 0 | 0 | 344 | 11 | 274 |
| 3 | 956 | 0 | 0 | 0 | 580 | 27 | 349 |
| 4 | 1837 | 0 | 83 | 0 | 1287 | 73 | 394 |
| 5 | 2987 | 1 | 806 | 1 | 1460 | 241 | 478 |
| 6 | 5345 | 2 | 2455 | 10 | 1476 | 525 | 877 |

Jackhmmer searches were seeded with three proteins A: TMEM106B, B: Vac7 (C-terminus only) C: Tag1. Hits were counted on the iterations indicated with domain assignments. Squares are coloured as follows: red – first occurrence; yellow – sharp increase (by factor of 5 or net increase ≥500); grey – moderate increase; blue – no or slow (≤20%) increase. Asterisk for “none” in C. indicates that Tag1 itself is not recognised to contain any domains by HMMER.

**Supplementary Table 4: Pairwise comparison of LEA-2 structures**

| | | | Z scores (RMSD / residues aligned) | | | | | |
|---|---|---|---|---|---|---|---|---|
| Structure | MSA seqs | TM-score (confidence) | 1YYC | LEA2-C (3BUT)* | LEA2-N* | T106B* | Vac7* | Tag1-C* |
| Plant LEA-2 (1YYC) | n.a. | | 23.9 (0/125) | 14.6 (2.5/125) | 11.6 (3.6/120) | 11.6 (3.0/122) | 9.1 (3.5/117) | 7.8 (4.5/118) |
| Archaeal LEA-2C* | 4889 | 0.811 (v.high) | | 26.6 (0.0/138) | 16.0 (2.1/127) | 13.0 (2.9/125) | 9.4 (2.8/114) | 9.0 (3.5/123) |
| Archaeal LEA-2N* | 10720 | | | | 26.1 (0/131) | 13.4 (3.0/123) | 11.5 (2.2/116) | 10.2 (2.7/121) |
| TMEM106B* | 3549 | 0.796 (v.high) | | | | 28.2 (0/157) | 10.5 (2.9/120) | 9.7 (3.5/126) |
| Vac7C | 1131 | 0.682 (high) | | | | | 32 (0/224) | 8.4 (3.5/123) |
| Tag1 – domain C | 483 | 0.838 (v.high) | | | | | | 28.2 (0/152) |

Pairwise structural comparisons were made using DALI for the plant LEA-2 domain NMR structures 1YYC, and predicted structures made by trRosetta without using template based information (indicated by asterisks) for both archaeal LEA-2 domains from *T. litoralis* WP_148290494.1 (C-terminus of which is highly related to 3BUT), TMEM106B, Vac7-C-terminus and domain C from Tag1.[38] The second column shows the number of sequences with at least 20 residues aligned for pairwise co-evolution, and the third column TM-score and confidence in each model. Scores in the final five columns are Z-scores for overall similarity from DALI, with RMSD and number of residues aligned in brackets.[39] Yellow background indicates strong alignment (Z-score ≥10). Squares on the long diagonal show self-alignments (grey background). "n.a." – not applicable.

**Supplementary Table 5. Annotations of PSI-BLAST hits to Vac7 in iterations 1 to 4**

| iteration | Significant hits (e.val.≤0.001) | | | Non-significant hits (e.val.≤1) | | |
|---|---|---|---|---|---|---|
| | Vac7 | LEA-2 | Other (no info) | Vac7 | LEA-2 | Other (no info) |
| 1 | 165 | 0 | 2 | 17 | 0 | 5 |
| 2 | 190 | 0 | 5 | 2 | 11 | 38 |
| 3 | 189 | 0 | 5 | 1 | 13 | 31 |
| 4 | 189 | 1 | 5 | 2 | 23 | 35 |

PSI-BLAST searches were seeded with the C-terminus of Vac7 (residues 932-1165) in the nr50 database, reporting hits with expected value ≤0.001 (significant) and above that but ≤1 (non-significant).
